# Methylphenidate exposure alters brain gene expression and induces transgenerational DNA methylation changes in *Poecilia reticulata* guppies

**DOI:** 10.64898/2025.12.16.694564

**Authors:** Rebekah J Alfaro, Alex R De Serrano, Dustin Sokolowski, Kimberly A Hughes, Helen Rodd, Ina Anreiter

## Abstract

Chronic exposure to stimulants is known to affect behavioral phenotypes and epigenetic profiles intergenerationally. We have previously shown that chronic methylphenidate hydrochloride (MPH) exposure in male guppies (*Poecilia reticulata*) induces persistent, paternally transmitted behavioral changes across multiple unexposed generations. Here, we investigated the underlying epigenetic signatures of this transgenerational behavioral inheritance. Building on our previous study, which used composite behavioral scores, we investigates the transgenerational inheritance patters of individual behaviors and found a robust, male-lineage–dependent increase in swimming across unexposed G2–G4 female offspring. Our molecular analysis showed that that chronic MPH exposure in G1 males alters brain gene expression, with 76 differentially expressed genes including genes with developmental and locomotory functions, and an over-representation of long non-coding RNAs. Furthermore, we found that unexposed G4 descendants from MPH-treated lineages exhibit widespread changes in brain DNA methylation, including genes related to developmental and swimming behavior. Interestingly, we found one lnRNA, LOC103476631, as both differentially expressed in G1 males and differentially methylated in G4 fish, providing a compelling candidate for ncRNA-directed DNA methylation as a mechanism for the observed transgenerational epigenetic inheritance of swimming behavior.

**Author Summary:** Drugs used to treat attention-deficit/hyperactivity disorder, such as methylphenidate, are widely prescribed to children and adolescents during key stages of brain development. We previously showed that exposing male guppies to a low, chronic dose of methylphenidate changed anxiety-like behavior not only in the treated fish but also in several generations of their unexposed descendants. In this study, we asked how such long-lasting behavioral effects might be recorded in the brain. We first confirmed that a simple measure of swimming behavior remains altered in descendants of treated males across multiple generations. We then examined brains from the directly exposed fathers and from great-grand-offspring that were never exposed to the drug. In fathers, we found changes in the activity of genes, including many long non-coding RNAs, a class of genes that has been linked to transgenerational inheritance. In great-grand-offspring, we found significantly altered DNA methylation profiles, an epigenetic mark that has also been associated with transgenerational inheritance. Together, our findings suggest that developmental exposure to a commonly used stimulant can leave a stable molecular ‘memory’ in the brain that persists across generations.

## Introduction

Behavioral traits arise from a complex interplay between genetic inheritance and environmental experience. Increasingly, research in epigenetics has revealed that environmental exposures can leave molecular imprints, the effects of which extend beyond the directly exposed individual, affecting offspring across multiple generations [1]. These findings challenge long-held assumptions that heritable traits are dictated solely by DNA sequence, and instead support a more dynamic view of inheritance, one in which experience, including early-life exposure to environmental stimuli or chemicals, can influence the biological and behavioral profiles of future generations [1]. Among the epigenetic mechanisms proposed to play a role in this transmission, DNA methylation has emerged as a central candidate, implicated in both stable gene regulation and non-Mendelian inheritance.

DNA methylation involves the addition of a methyl group to CpG base pairs and has been shown to regulate gene expression without altering the underlying DNA sequence [2]. This mechanism underlies classic examples of transgenerational epigenetic inheritance (TEI) in mammals, including coat color variation in mice [3], maternal grooming behavior in rats [4], and diet-induced phenotypic changes in livestock and rodents [5–7]. Other studies have also shown that environmental experiences, such as trauma, can induce lasting changes to DNA methylation that are passed from parent to offspring [8]. Furthermore, exposure to chemicals, including endocrine disruptors, pesticides, and simulant drugs can induce epigenetic modifications that persist across generations. For example, in rodents, exposure to vinclozolin, a fungicide, results in transgenerationally inherited changes in sperm DNA methylation [9,10], prenatal paternal nicotine exposure results in heightened fear responses in the unexposed F1 and F2 generations and is associated with increased methylation of several genes [11], and chronic paternal cocaine use increases addiction susceptibility in F1 and F2 generations, correlating with persistent changes in sperm DNA methylation [12]. However, despite the growing number of environmental triggers investigated, our understanding of the mechanisms, consistency, and specificity of TEI remains incomplete.

One particularly understudied class of chemicals in this context are medically prescribed stimulants, such as methylphenidate hydrochloride (MPH; commonly known under the brand name Ritalin). MPH is a widely prescribed medication for attention-deficit/hyperactivity disorder (ADHD) in children, adolescents, and adults [13–15], yet its long-term neurodevelopmental and transgenerational effects in humans remain largely unexplored. In rodents, chronic MPH administration can alter reward sensitivity, exploratory behavior, and gene expression related to neuroplasticity [16–18]. It has also been associated with effects on testicular morphology and sperm development [19]. As far as we are aware there are no published studies examining the epigenetic consequences of MPH exposure beyond the directly treated individual and its immediate offspring.

Given the parallels in the molecular mechanisms of action between MPH and other stimulants shown to induce TEI [20], and the well-documented neurobehavioral effects of MPH, the absence of studies investigating its transgenerational potential represents a critical gap. Addressing this gap is particularly important in the context of widespread MPH use among youth during sensitive periods of neurodevelopment.

In this study, we leverage a vertebrate model, the Trinidadian guppy (*Poecilia reticulata*), to investigate a potential epigenetic mechanism for the effects of chronic low-dose MPH exposure in one generation on behavioral alterations in subsequent, unexposed generations. Teleost fish share several key features of neurodevelopment with mammals [21], and poeciliids exhibit mammal-like patterns of DNA methylation [22, 23]. As live bearers, guppy embryos develop *in utero*, allowing for possible *in utero* transmission of molecular changes, a key aspect of TEI in mammals [1]. A previous study found that exposure of juvenile guppies (G1) to MPH induced anxiety-like behaviors in adulthood, and that these behavioral differences persisted in the G2 through G4 generations despite no direct exposure in those later generations. This transgenerational transmission occurred via the paternal lineage only (when a male G1 fish was exposed to MPH).

Building on this work, we investigated the molecular underpinnings of the transgenerational transmission of MPH-induced behavioral differences. We first confirmed the persistence of behavioral alterations across generations in De Serrano *et al*. [24] at the levels of individual behaviors, and subsequently examined gene expression changes in the directly exposed G1 generation, and assessed genome-wide DNA methylation patterns in the unexposed G4 generation. We found that the transgenerational behavioral effects of MPH were associated with differential gene expression patterns in the treated fish and intergenerational changes in DNA methylation, and that these changes were detectable in genes associated with development and behavior. By integrating behavioral, transcriptomic, and epigenomic analyses, this work provides novel insights into the potential for medically prescribed stimulants to elicit long-term and heritable biological changes.

## Results

### Behavior

We assessed two individual behaviors that were most frequently performed by fish in the open field arena across all four generations: total time spent swimming in the tub (Swim) and number of outer squares traversed (outer squares) (Fig 1). We focused on individual behaviors because we thought we would be more likely to detect clear associations with gene expression and methylation than with the complex, composite behavioral metrics in De Serrano *et al*. [24]. We asked if there were significant, direct effects of MPH on these behaviors of G1 fish, if there were effects of G1 MPH treatment on G2-G4 fish, and if any of those effects of MPH were consistent across the G2-G4 generations. Generally, consistent with those of De Serrano *et al*. [24], our results showed that there was a significant effect of male G1 treatment alone or in an interaction on individual behaviors of the fish in G2-G4 generations and that the effects of transgenerational MPH were similar across those generations. However, in contrast to De Serrano *et al*. [24], who observed a significant sex* male G1 treatment interaction for G1fish, in this study, for the duration of time spent swimming (Swim), there was a significant effect of sex (Fig 1c, d, Table 1) but not a significant effect of treatment alone or interactions with the other factors we assessed.

**Fig 1:**
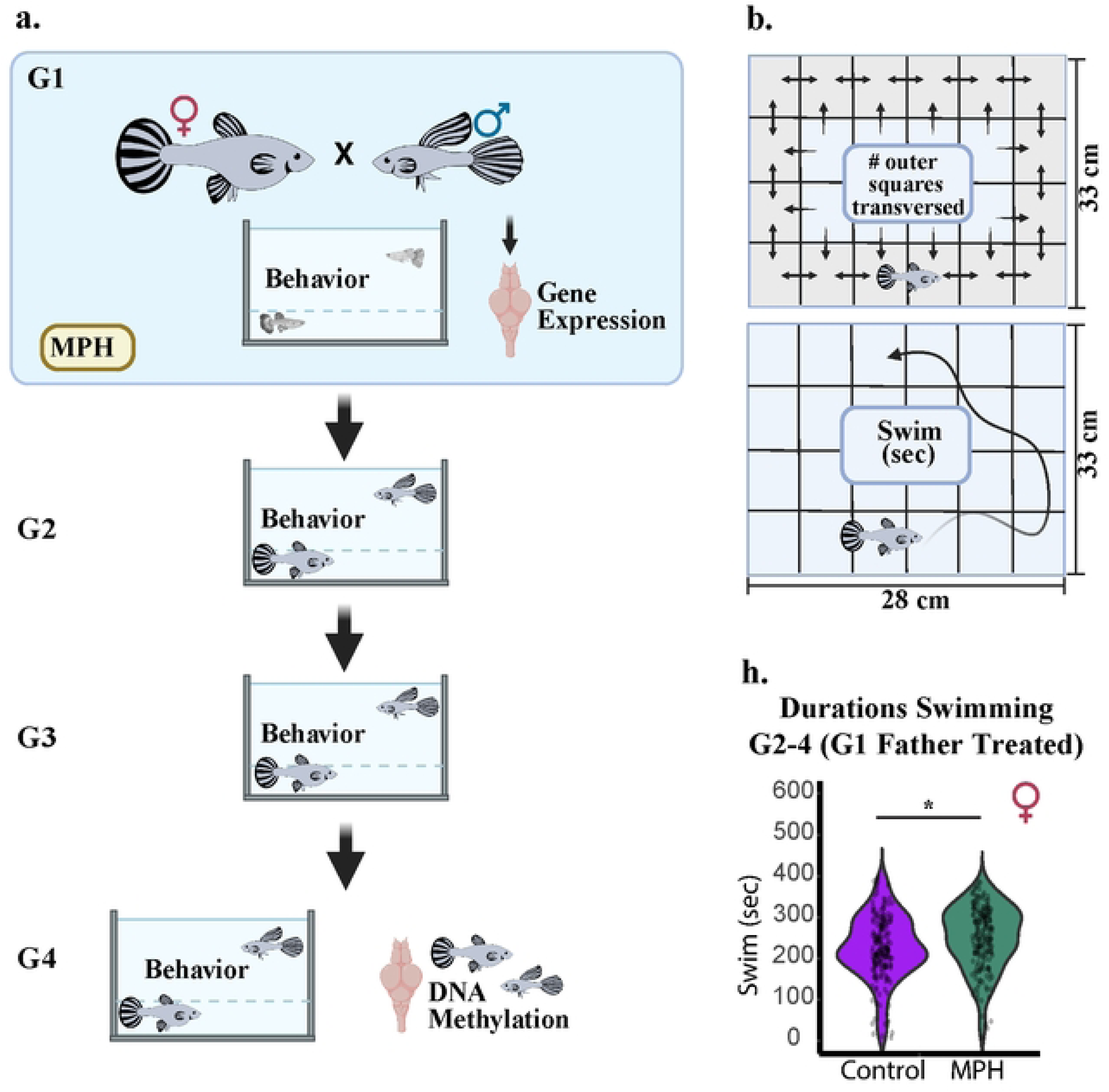
Behavior in open field trials. **a.** Experimental design: G1 fish were treated with MPH or water (control), gene expression was assessed in the brains of G1 males, behaviors were assessed every generation (G1-G4), and DNA methylation was assessed in the brains of G4 fish. **b.** Individual behaviors assessed in this study: outer squares (number of outer squares entered in the open field test) and Swim (time spent swimming in the open field test). **c.** For G_2_-G_4_ females, having a male G_1_ ancestor that was treated with MPH had a significant effect Swim scores (t=−3.17, df=358, p=0.0017), with MPH-treated G_1_ male ancestors lead to more time spent swimming. * p < 0.05, ** p < 0.01, *** p < 0.01

**Table 1:**
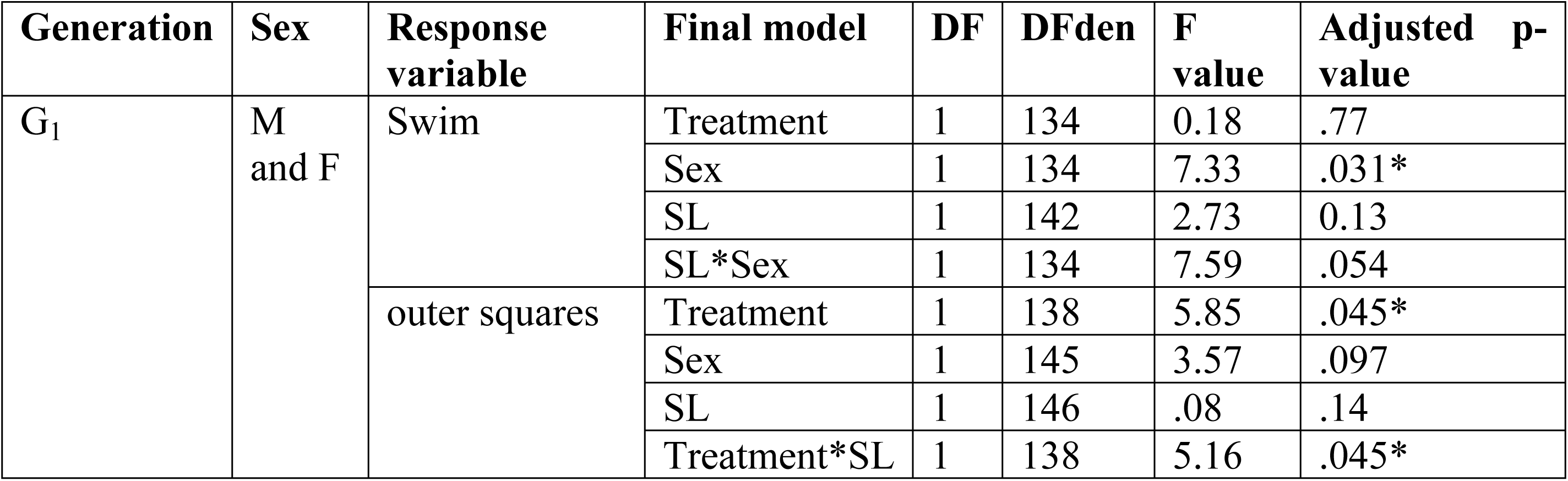
Mixed model analyses of behavior scores for G_1_ Swim and outer squares. Covariates are included where they were significant in the final analysis. * p < 0.05, ** p < 0.01, *** p < 0.01

In this study, for G2-G4 females, for the duration swimming, there was not a significant effect of generation, either alone or in an interaction, but there was a significant effect of treatment received by G1 male ancestors (Fig 1h, Table 2) where G2-G4 females with an MPH-treated male G1 ancestor (females from the MPH and CTR/MPH lineages) spent significantly more time swimming during the open field test than females with a control G1 male ancestor (females from the CTR and MPH/CTR lineages). For G2-G4 males, there was not a significant the effect of G1 ancestor treatment on duration swimming although the pattern for the effects of male G1 treatment was in the same direction as for G2-G4 females (Fig 1g, Table 2). Comparing the descendants of MPH-treated and control fish across generations, for every generation (G2-G4), females with an MPH-treated male ancestor spent significantly more time swimming than females with control ancestors and there was not a significant generation*treatment interaction so the extent of the differences between treatments were similar across the three generations (Fig S1; Table 2). There was not a significant effect of the ancestors’ treatments on male G_2_-G_4_ Swim scores, nor was there a significant interaction between generation and G_1_ treatment, so we could not address the question of whether the effects of MPH treatment were consistent across generations. However, for the G_2_-G_4_ males there was a significant generation*age at testing interaction (Table 2).

**Table 2:**
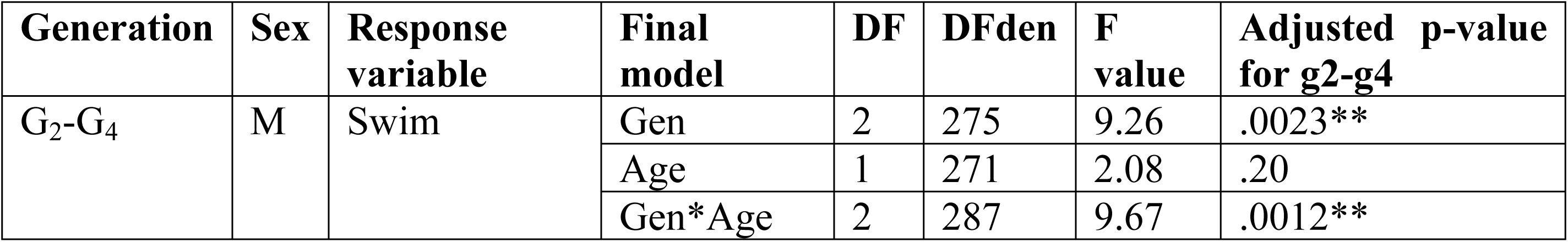
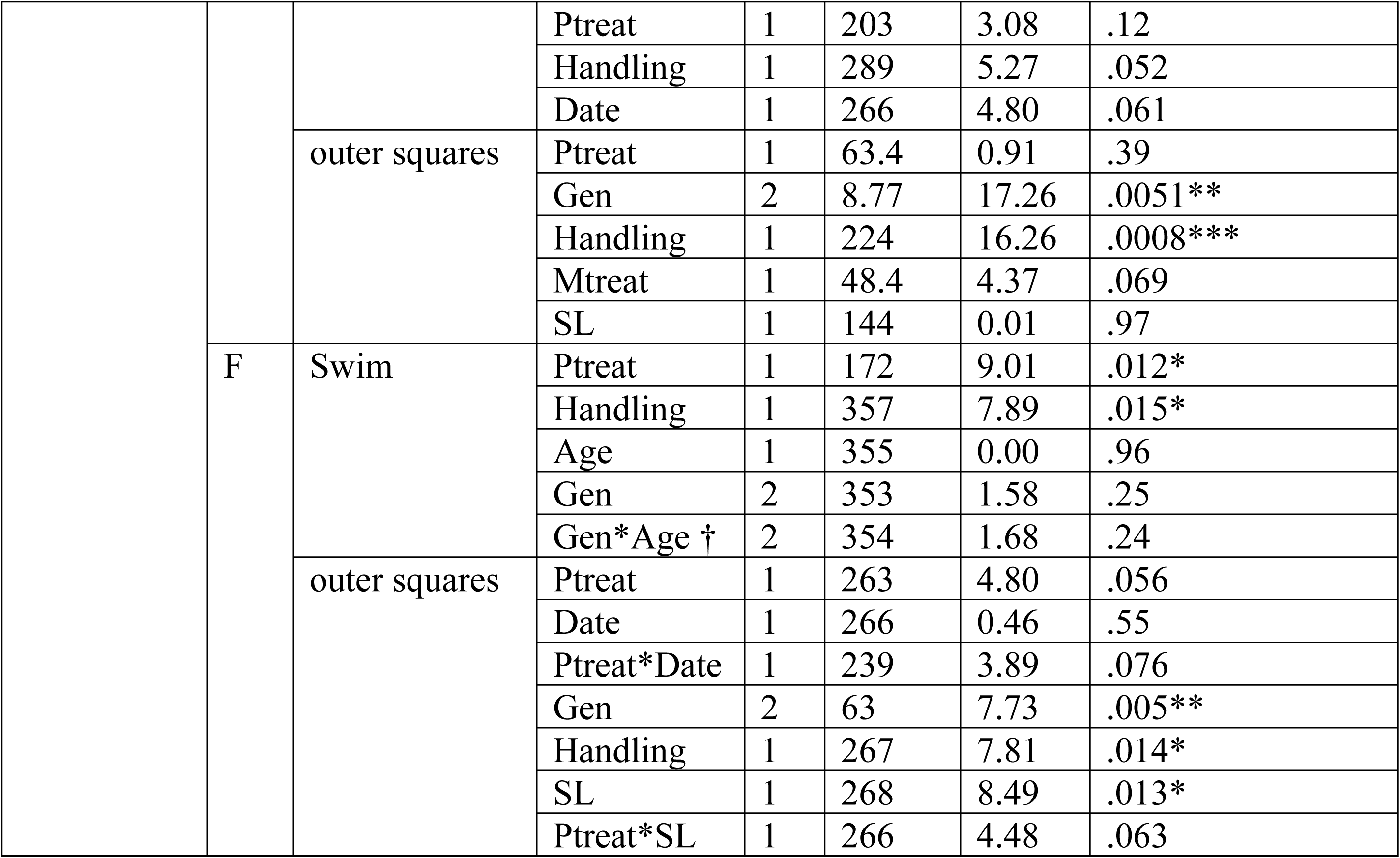
Mixed model analyses of behavior scores for Swim and outer squares. Sexes were analyzed separately in G2-G4 because standard length was measured differently for males and females in these generations. Ptreat is the treatment that the male ancestor in G1 received; Mtreat is the treatment that the female ancestor in G1 received. Date is the date the fish was tested and euthanized. Covariates are included where they were significant in the final analyses. * p < 0.05, ** p < 0.01, *** p < 0.01

For the G1 fish, for outer squares, there was a significant interaction between MPH and standard length for MPH-treated guppies such that the number of squares decreased with SL (F_1,63.3_=8.62, p=.005) but that relationship was not significant for control fish (F_1.6.75_=.08, p=.78), (Table 1). Treatment did not interact significantly with any of the other variables.

For G2-G4, for outer squares, there were significant effects, but no significant effect of treatment. Both males and females in G2-G4, for the number of outer squares entered (Table 2) there was not a significant effect of G1 ancestor treatment either alone or in an interaction for either sex. However, for females, the treatment of the male G1 ancestor showed a trend but did not reach statistical significance (p=0.056), with controls entering fewer outer squares than treated fish. This trend was not a result of the effect of treatment on overall activity levels, which was evaluated when the fish were in their home tanks [24]. Generation had a significant effect on outer squares by both males and females in G2-G4 with G2 males having significantly higher outer squares than G3 and G4 fish (Fig S1; Table 2); for females, G4 individuals showed the highest outer squares compared to G2 and G3 females (Fig S1; Table 2). Handling time also significantly affected the outer squares scores for both sexes (Table 2).

Together, our results, in alignment with those of De Serrano *et al*. [24], allowed us to proceed with the main goal of this study to look for associations between the effects of male G1 MPH treatment on behavior, gene expression patterns, and offspring brain methylation.

### MPH affects gene expression profiles in G1 males

To explore MPH-induced changes in gene expression in G1 fish, we performed RNA-seq on brains of G1 males. We focused on males because composite behavioral effects were transmitted through the paternal line [24]. Principal component analysis of G1 male brain expression profiles showed greater within-group variability in the control group than the treated group (Fig S2). We identified 76 differentially expressed genes (DEGs; Supplemental data file 1), of which 61 were upregulated and 15 were downregulated in the treated fish relative to the control fish (Fig 2 a-c). Fifty-three of these DEGs were protein coding genes (Supplemental data file 1), while 22 were non-coding, and one was unannotated. The DEG analysis accounts for the large number of genes tested using Benjamini-Hochberg False Discovery Rate (FDR) correction, which is implemented by default in the edgeR::topTags() function. In total, 21,909 genes were tested. At an FDR cutoff of 0.05, we would expect approximately 0.05×21,909=1,095.45 genes to be significant by chance alone, assuming a null distribution with no true differences. However, our filtering strategy further included a biological significance threshold (|logFC| > 1.5), which reduces the likelihood that these 76 DEGs are false positives. These findings show a significant over-representation of non-coding genes in our list of DEGs (Fisher’s exact test Odds ratio = 6.4014, p-value = 4.41e^−10^) (see Supplemental data file 1 for number of reads per sample). Among the most significant DEGs were genes with potential links to swimming behavior: dynein-coding gene (*dnah3)*, two hox genes (*hoxc8a*, *hoxb6-like*), and the long non-coding RNAs *LOC103476631* and *LOC108167276* (Fig 2 d-h). We did not find any GO terms or pathways significantly enriched in our list of DEGs. We included annotated functions (GO terms) for significant protein coding DEGs in Supplemental data file 1.

**Fig 2:**
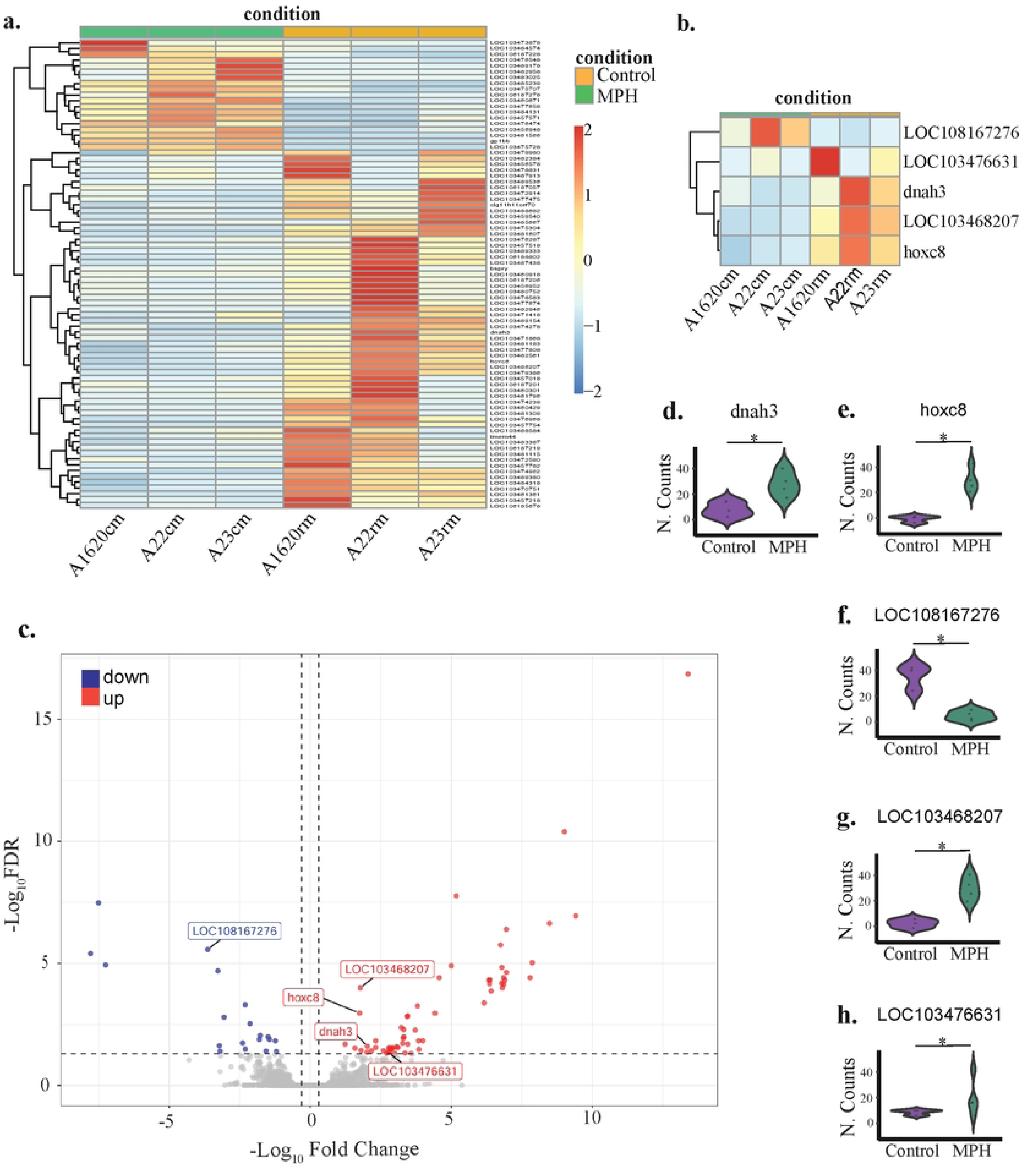
Gene expression profiles in MPH treated and control G1 males. **a.** Heatmap of all differentially expressed genes between control and MPH treated male brains. **b.** Heatmap of all differentially expressed genes of interest between control and MPH treated male brains. **c.** Volcano plot of the 76 genes that were differentially expressed between control and MPH treated male brains. Differentially expressed genes of interest are labeled. **d-h.** Normalized count plots for differentially expressed genes of interest. *Hox-b6a-like* (LOC103468207; p= 9.98e^−5^), *Hoxc8a* (p= 0.001084)*, Dnah3* (p= 0.024777), and *LOC103476631* (p= 0.045869316) are upregulated in MPH-treated fish, LOC108167276 (p= 2.71e^−6^) is downregulated in MPH-treated fish. * p < 0.05, ** p < 0.01, *** p < 0.01.

### MPH treatment affects DNA methylation patterns of G4 progeny

Given our observations on the effects of MPH treatment on the transgenerational effects on behavior, and the differential gene expression patterns in the brains, we next used bisulfite sequencing to find epigenetic signatures in G4 progeny that might explain the transmission of behavior. As with G1 gene expression profiles, principal component analysis of G4 brain methylation profiles showed greater within-group variability in the control group than the treated group (Fig S2). We identified 3026 differentially methylated cytosines (DMCs; Supplemental data file 2), of which 58% (1753 DMCs) were hypermethylated and 42% (1273 DMCs) were hypomethylated (Fig 3a). When examining the genomic context of the DMCs on annotated chromosomes and genes (2653 out of 3026 DMCs), most were in exons (42%), while 10% were in promoters (10%) (Fig 3b). DMCs in introns (%) and intergenic regions (%) made up the rest of the DMCs. Functional profiling of all annotated DMCs yielded only one significantly enriched GO term in our set of DMCs: anatomical structure development (Biological Process GO:0048856; p=4.470e-4). There were no significantly enriched pathways. However, several Human Phenotype Ontology terms were significantly enriched: phenotypic abnormality (HP:0000118; p=3.543e-3), abnormal appendicular skeleton morphology (HP:0011844; 1.709e-2), abnormality of the nervous system (HP:0000707; p=4.419e-2), and abnormality of the upper limb (HP:0002817; p=4.656e-2).

**Fig 3:**
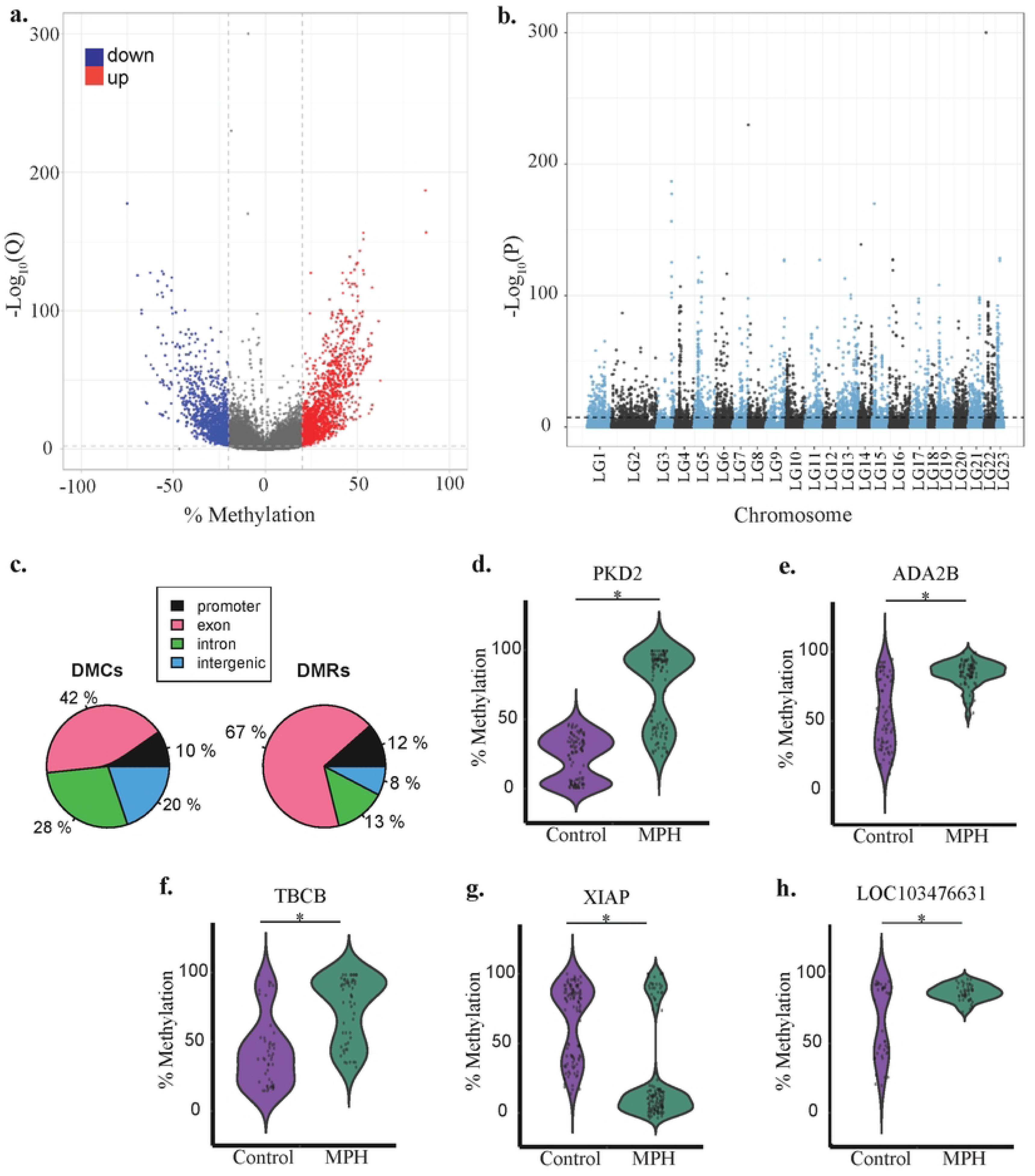
DNA methylation profiles in G4 offspring. **a.** Volcano plot for 550,352 DMCs in the brains of G4 fish. **b.** Distribution of DMCs along the 23 chromosomes of the Guppy genome. **c.** Genomic contexts of differentially methylated cytosines (DMCs) and differentially methylated regions (DMRs) When classifying the genomic context of a DMR, precedence was given to promotors > exons > introns > intergenic regions when a DMR spanned multiple structures. **d-h.** Percent methylation differences between G_4_ descendants of MPH-treated G_1_ fish (n=9) and G_4_ descendants of control G_1_ fish (n=8) for five differentially methylated regions of interest. DMRs displayed were within 2000bp of the TSS and located on the promotor region or on the first two exons. The DMRs at *pkd2-l1, ada2b, LOC10347663, tbcb* had significantly higher mean percent methylation in MPH fish compared to control fish (40.95%, 42.65%, 36.22%, 38.25% respectively). The DMR at *xiap* had a significantly lower mean percent methylation in MPH fish compared to control fish (−39.68%). Statistical significance was assessed using Wilcoxon rank-sum tests with the following p-values: pkd2-l1: 2.61e⁻²⁷, ada2b: 2.34e⁻¹⁴, xiap: 6.84e⁻¹⁸, tbcb: 4.10e⁻⁸, LOC10347663: 2.08e⁻².

To test if gene function was affected by differential methylation, we assessed whether differentially methylated cytosines (DMCs) clustered in specific genomic regions. We identified 58 differentially methylated regions (DMRs, Supplemental data file 2); 52 of these DMRS were located on annotated chromosomes and the remaining 6 DMRs were located on unplaced scaffolds. Like the DMCs, most of the annotated DMRs were in exons (67%) with the remaining DMRs split between promoters, introns, and intergenic areas (Fig 3b). Thirteen of the DMRs were within 2000bp of the transcription start site (TSS) and located in a promoter or exon (Supplemental data file 2), and thus more likely to alter gene expression [25]. As with DEGs identified in G1 males, some genes that had DMRs along the downstream regions had annotated functions that may affect the observed swimming behavioral phenotypes and their transmission, such as *pkd2-l1*, *xiap*, *tbcb*, and *ada2b*, (Fig 3c-g). The MPH group tended to have a higher mean percent methylation at these DMRs, except for the DMR on *xiap*, in which the MPH group had a smaller mean percent methylation than the Control group (Fig 3f). The DMRs on *pkd2-l1*, *ada2b*, and *xiap* were the DMRs on annotated genes that had the largest differences in methylation between groups (Fig 3c-g; Supplemental data file 2; *pkd2-l1* q=4.35072e^−95^, *ada2b* q=6.8975e^−100^, *xiap* q=2.2451e^−99^). Notably, lnRNA, LOC103476631, which was upregulated in the brains of G1 males treated with MPH (Fig 2h; q=2.08e⁻²), was also among the most significantly differentially methylated genes in G4 progeny (Fig 3h).

### Changes in behavior are associated with Methylation at *pkd2-l1*

To see if methylation differences at the five genes we have identified as being potentially driving swimming behavior could explain the transgenerational differences in the behavior of G4, we looked for an association between % methylation at these loci and the behaviors we scored. For outer squares, only *pkd2-l1* had a significant (negative) association with percent methylation at its DMR (Fig 4; Table 3). There was significant interaction between treatment group and % methylation of *pkd2-l1*, such that the slope was steeper for MPH fish than for the CTR fish, where the slope was not significant. There was also a significant sex*treatment interaction for this behavior. For swim, there were no significant associations between % methylation at these loci or any two-way interactions between them and treatment (results not shown).

**Fig 4:**
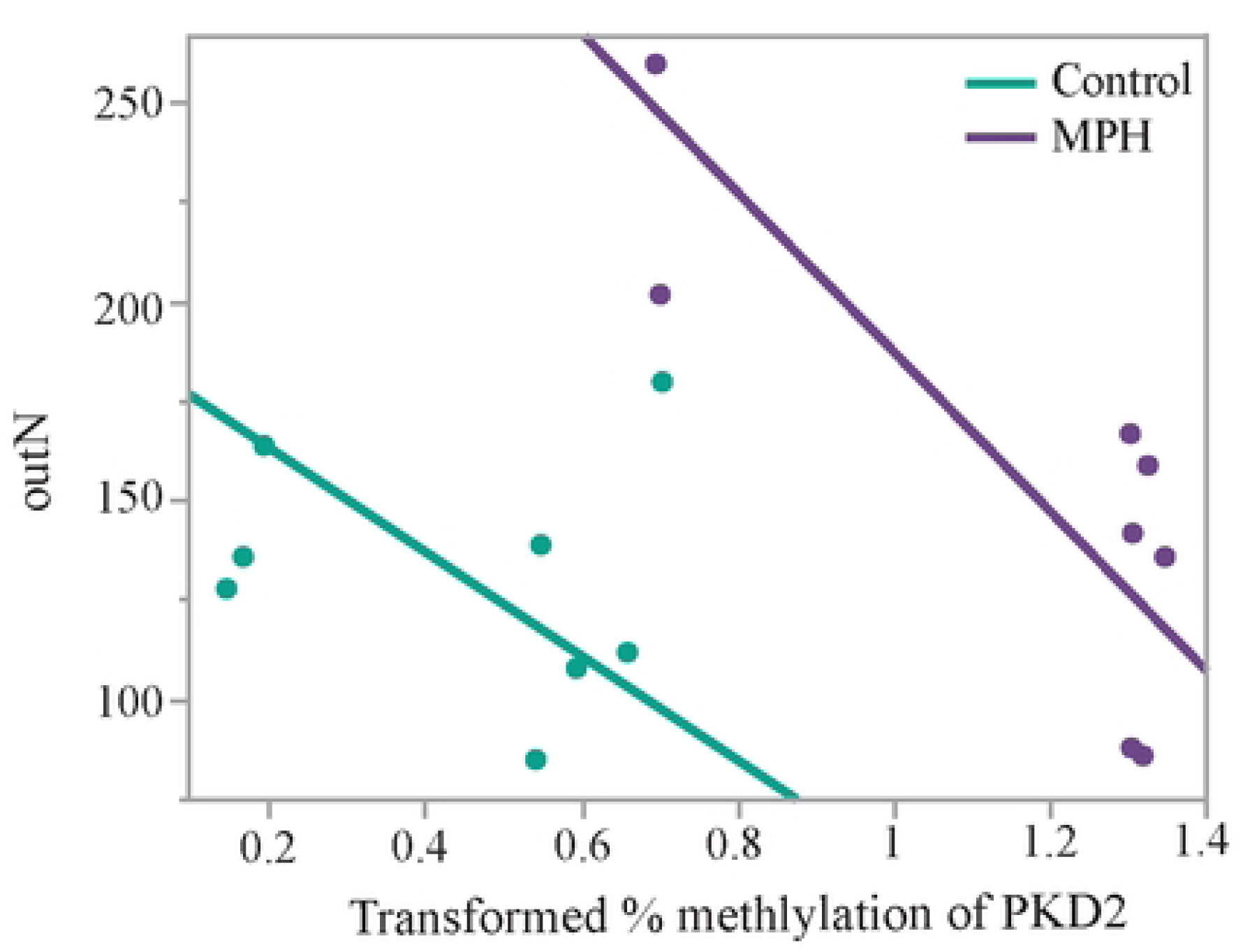
Association between percent methylation of *pkd2-l1*, and outer squares entered. As the mean percent methylation on the differentially methylated region of *pkd2-l1* increased, the number of outer squares entered during the open field test (outer squares) decreased. A reduced major axis regression analysis indicated that the slope is significant for G_4_ fish with MPH-treated G_1_ ancestors (red symbols, slope=−200.45, p<.025), but not for ones with control ancestors (blue symbols, slope=−130.54, p=0.74).

**Table 3:** Results of the final analysis of covariance (ANCOVA) of the response variable, outer squares entered during the open field test (outer squares), in relation to transformed mean % methylation for five genes (*pkd2-l1*, *xiap*, *tbcb*, *ada2b*, and *LOC103476631*) and treatment group along with sex. Mean % methylation refers to the transformed mean methylation percentage across the entire differentially methylated region located on that gene. Treatment refers to the treatment group of their G_1_ ancestors. P values for genes and interactions that were greater than 0.1 were removed from the final analysis. * p < 0.05, ** p < 0.01, *** p < 0.01

| Final model | Estimate | DF | DFden | F ratio | p-value |
| --- | --- | --- | --- | --- | --- |
| Intercept | 83.61 |  |  |  |  |
| Treatment | 10.38 | 1 | 9 | 0.12 | .740 |
| Sex | 7.02 | 1 | 9 | .90 | .367 |
| Treatment*Sex | -18.72 | 1 | 9 | 6.22 | .034* |
| Mean % Methylation pkd | -29.71 | 1 | 9 | 0.35 | .570 |
| Treatment*Mean % Methylation pkd | 147.72 | 1 | 9 | 11.18 | .009** |
| Mean % Methylation ada | 129.96 | 1 | 9 | 3.54 | .092 |

## Discussion

This study represents the first investigation into the potential epigenetic pathways that mediate transgenerational behavioral alterations in response to methylphenidate (MPH) exposure. Following up on our previous behavioral findings [24], we showed that chronic low-dose MPH exposure in one generation results in epigenetic alterations in subsequent, unexposed generations through sex-specific inheritance patterns with male G1 treatment affecting both G1 gene expression and methylation patterns in later generations. The identification of lncRNA LOC103476631 as both differentially expressed in G1 fish and differentially methylated in G4 fish provides compelling evidence for ncRNA-directed DNA methylation as a mechanism for transgenerational epigenetic inheritance. This aligns with recent evidence demonstrating that noncoding RNAs can circumvent germline reprogramming by being loaded into germ cells and facilitating the establishment, maintenance, and inheritance of epigenetic marks [26–28]. This finding is particularly significant given that similar mechanisms have been observed in rodent studies where paternal stress resulted in transgenerational behavioral patterns associated with differential expression of ncRNAs and de-novo establishment of tissue-specific differentially methylated regions across generations [29].

The transgenerational effects observed through the paternal lineage directly connect to previous findings that MPH affects testicular morphology and sperm development [19].The identification of pkd2-l1 as a key differentially methylated gene is particularly relevant in this context as it is expressed in testes and plays crucial roles in ciliary function and mechanosensory processes essential for sperm motility [30]. The significant association between pkd2-l1 methylation and behavioral outcomes provides a molecular link between sperm-mediated inheritance and phenotypic transmission. In addition, the involvement of genes related to microtubule dynamics (dnah3, tbcb) further supports the connection to sperm development, as proper microtubule function is essential for sperm flagellar movement and overall male fertility. Pkd2-l1 proteins are expressed by cerebrospinal fluid-contacting neurons (CSF-cNs), which monitor CSF composition and flow [31]. This system is conserved in the spinal cord across bony vertebrates [32] and is essential for ciliary function [33]. Further supporting the role of pkd2-l1 in modulating locomotor activity, adult pkd2-l1 mutant zebrafish displayed an exaggerated rounding of the spinal column that arose spontaneously during development and reduced their tailbeat frequency [34], this phenotype was more severe in males [35]. It is thus possible that both the paternal transgenerational transmission patterns, and the effects on swim in our open field test are mediated by altered ciliary function to some extent.

Another compelling potential mediator for TEI in our study involves the differentially methylated region on ada2b (adenosine deaminase 2B), which had the second-largest methylation difference between treatment groups. Ada2b represents a potentially novel regulatory axis affecting swimming behavior through adenosine signaling pathways. The methylation changes at ada2b could alter extracellular adenosine metabolism [36], Adenosine signaling through adenosine A2A receptors (A2AR) play crucial roles in both sperm function and swimming motor control [37]. A2AR is expressed both in sperm tails, where its levels positively correlate with progressive motility [38], and in the central nervous system, where it affects swimming behavior [39]. Furthermore, A2ARs are G protein-coupled receptors (GPCRs) that modulate dopamine signaling, potentially modulating the effects of MPH, which primarily binds to dopamine receptors [39]. Further supporting this role of regulation by gpr148 (G protein-coupled receptor 148) was differentially methylated in the G4 in our study. While the ligand of gpr148 is currently not known, this receptor is primarily expressed in the nervous system and testis and has been linked to ADHD in humans [40]. Thus, Inosine and GPCR signaling presents a second plausible avenue for both the paternal transgenerational transmission patterns, and the effects on fish behavior in a novel environment that we see in our study.

An additional observation in our data is the reduced within-group variability in both RNA-seq expression profiles and DNA methylation patterns in MPH-treated samples compared to controls. The reduced within-group variability suggests that MPH treatment induces a convergence toward a more similar molecular setpoint, potentially representing reduced genomic entropy (a measure of randomness in the variation in gene expression) following drug exposure. This molecular convergence may represent an adaptive response to MPH that becomes heritable through epigenetic mechanisms, ensuring consistent transmission of beneficial regulatory changes across generations. This would align with evidence that methylphenidate can restore better synchronization between brain networks and reduce moment-to-moment variability in ADHD patients [41].

This study establishes that MPH is capable of inducing transgenerational epigenetic inheritance through mechanisms potentially involving lncRNA-directed DNA methylation and convergence toward reduced genomic entropy. The identification of sex-specific inheritance patterns and the possible connection to sperm development provides a mechanistic framework for understanding how therapeutic stimulant use can have lasting effects across generations. This has implications for human health given the widespread use of MPH among youth during sensitive periods of neurodevelopment [42]. As such, transgenerational inheritance of drug-induced modifications warrants careful consideration, and future studies should examine other epigenetic mechanisms, including histone modifications and chromatin remodeling, which may contribute to the transgenerational effects observed, as well as investigate the temporal stability of these epigenetic modifications across additional generations and their potential reversibility.

## Methods

### Experimental design

This study used the same fish as our previous behavioral study [24]. Briefly, a reciprocal treatment design with G1 mothers, G1 fathers, or both being exposed to either MPH dissolved in water or water alone (control), as described in Briefly, G1 individuals were treated chronically with either MPH (group MPH) or water (group CTR). The G1 were mated using a factorial design to produce four offspring treatment groups in the G2 that were maintained to produce the G3 and G4. The G2-G4 were never directly treated with MPH and their division into four treatment groups in each generation is based on the treatment that their G1 ancestors received. The four groups in each generation in the G2-G4 are: MPH (both G1 ancestors treated with MPH), CTR/MPH (male G1 ancestor treated with MPH mated with a control female), MPH/CTR (female ancestor treated with MPH mated with a control male), and CTR (both G1 ancestors were controls).

### Behavior

We investigated the inheritance patterns of the two most frequently performed individual behaviors in the open field test, Swim, the duration of time spent swimming during the open field test, and outer squares, the number of outer squares of the arena entered during the open field test. Both variables were included in the composite behavioral scores in De Serrano *et al*. [24]. Analyses for G_1_ fish were done separately from G_2_-G_4_, as they were the only generation treated directly with MPH. Sexes were analyzed separately for G_2_-G_4_ because standard length (SL, measure of body size), which can contribute to variation in behavior in open field tests, was measured differently for males and females, and sex was a significant covariate in some analyses. We used a linear mixed model, using the Proc Mixed function, in SAS v9.4. All of the models for G_2_-G_4_ fish included G_1_ male treatment, G_1_ female treatment, age at testing, date tested and euthanized (fish were euthanized on the date that they were tested), generation, standard length, handling time (time spent catching and moving fish from their tank to the open field tub with a net), and all two and three-way interactions as fixed effects. Pedigree (the relatedness among individuals) was included as a random effect. Any covariates and interactions where p > 0.1 were removed in a stepwise fashion from the final model. Behavior data for the G_1_ was analyzed in the same way, where treatment, age at testing, sex, date tested, time spent handling the fish, SL, and their interactions were included as fixed effects in the original model. Sibling ID (what sibling group they belonged to) was included as a random effect. All data met the assumptions of the analysis. Least-squares mean behavior scores were obtained from the final models using the SAS lsmeans function and post-hoc comparisons were performed using the simulate function. To control for the risk of false positives when comparing across the sexes and because we were analyzing two related behaviors, we adjusted the P-values by false discovery rate (FDR) using the Benjamini-Hochberg procedure. Violin plots were made using ggplot2 v.3.5.1 [42] in R v4.3.2 for Windows, the version of R used for all subsequent analyses.

### Sample Collection

All focal guppies were sexually mature and were euthanized directly following behavioral testing by placing them in an ice slurry for 10 seconds and decapitating them using a scalpel. Whole brains were collected by cutting the decapitated head down the center line and removing the brain tissue, which was placed in microcentrifuge tubes with RNAlater and stored at −20° C [24]. DNA and RNA from whole brain tissue was extracted using the Qiagen DNeasy Blood & Tissue Kit (Qiagen, Hilden, Germany) and the RNeasy Lipid Tissue Mini Kit (Qiagen, Hilden, Germany).

### RNA Sequencing

Pooled RNA-seq data were collected from four MPH-treated (R) and three control (C) males from the parental generation (G1). We focused on males because composite behavioral differences were transmitted via the paternal line in [24]. All pooled RNA samples consisted of RNA from two male siblings in the same treatment group, except for one sample each for both the control and MPH-treated groups, where RNA from two male siblings and one unrelated male individual from the same treatment group were pooled. Library preparation, including cDNA synthesis, was performed using the NEBNext Ultra RNA Library Prep Kit for Illumina (NEB #E7530) according to the manufacturer’s instructions. Libraries were sequenced on an Illumina HiSeq 2500 instrument (Illumina Inc., San Diego, CA) at Florida State University, running HCS v2.2.38 software. Quality controls for sequenced reads were performed using FastQC v0.11.2 [44] and adapter removal was performed using Trimmomatic v0.32 [45]. Cleaned reads were aligned to the reference genome (GCF_000633615.1) using HISAT2 v2.2.1 [46] and gene expression was estimated using StringTie v2.2.1 [47]. We filtered genes with low expression by removing genes with a mean read count ≤ 5 across all samples, then further filtered genes to retain only those expressed (with read counts > 5) in at least two samples. We also applied group-specific filters requiring presence in at least 2–3 samples per group depending on the group. The R package edgeR v4.0.16 [48] was used to identify genes that were differentially expressed (DEGs) between two groups. A gene was defined as a DEG if it had a minimum of a 1.5-fold change in abundance and FDR < 0.05 for pairwise comparisons between the two groups. The R package RUVSeq v1.36.0 [49] was used to aid in normalization of the raw counts that were input into edgeR.

The basic R stats package v3.6.2 was used to build a custom script to identify whether DEGs are protein coding or noncoding using version GCF_000633615.1 of the annotated guppy genome. The fisher.test function was used to test whether DEGs were enriched for noncoding genes. DEGs were input into g:Profiler to obtain a list of enriched GO terms, KEGG pathways, and Human Phenotype Ontology (HP) terms [50]. A statistical domain scope that included only annotated genes and a significance threshold of p<0.05 with a g:SCS threshold for multiple testing correction was used in the functional profiling. Single gene plots of DEGs were built using a custom script with functions from the R package ggplot2 [43].

### DNA Methylation

Reduced representation bisulfite sequencing (RRBS) data was collected for whole brains of nine G4 fish (6 females, 3 males) from the MPH-treated lineage (MPH) and eight G4 fish (5 females, 3 males) from the control lineage (CTR). The fish were of different ages when they were collected, and they were collected at different times of year (see Supplemental data file 2). Library preparation, bisulfite conversion and amplification were performed by Diagenode Inc. (RRBS Service, Diagenode Cat# G02020000) using their Premium RRBS Kit (Diagenode Cat# C02030033). Libraries were sequenced on an Illumina NovaSeq6000 instrument (Illumina Inc., San Diego, CA) in two batches; the first running NovaSeq Control Software v1.6.0 for the 2022 (first batch) samples and v.1.7.5 for the 2024 (second batch) samples. RTA v3.4.4 and bcl2fastq 2.20 v2.20.0.422 were used for the 2024 samples during the sequencing process.

Quality control was performed using FastQC v0.11.8 [44] and adapter removal was performed using Trim Galore! v0.4.1 [51]. Cleaned reads were aligned to the reference genome (GCF_000633615.1) using Bismark v0.16.1 for the 2022 samples and v0.20.0 for the 2024 samples [52]. More specifically, the cytosine2coverage and bismark_methylation_extractor modules of Bismark were used to infer the methylation state of all cytosines for every single uniquely mappable read, to infer their context, and to compute the percentage methylation. The resulting reported cytosines were filtered so that only the CpGs covered in each sample remained. The data shows an enrichment of small p values, providing good support for a class of CpGs that significantly respond to ancestral MPH treatment (Fig S3) [53].

Analyses of differential methylation comparing MPH and CTR samples were conducted using the R package Methylkit v1.29.1 [54]. To be considered statistically significant, differentially methylated cytosines (DMCs) had to have a q-value cutoff < 0.01 and a methylation difference greater than 20%. Age at testing, sequencing batch, date of euthanasia, and sex were included as covariates in the computation of differential methylation. Using the DMCs identified by Methylkit, DMRs (differentially methylated regions) were identified using the R package eDMR version 0.6.4.1 [55] with default parameters. To be considered significant, a DMR had to contain at least three CpG sites within an algorithm-specified genomic distance, with at least one classified as a DMC.

The R package genomation v1.34.0 [56] was used to identify the genomic context of the significant DMRs, as in Hu *et al* [22]. The proximity of each DMR to the nearest known transcription start site (TSS) was annotated and the DMRs were classified into four groups, depending on the region of the gene in which they were located: exons, introns, promotors, and intergenic regions. If a DMR spanned multiple regions, precedence was given to promoters > exons > introns > intergenic regions when classifying the genomic context of the DMR [22]. Genomic context could only be evaluated for the DMCs that were on genes with annotations, and unannotated genes were excluded from this analysis. Methylation data were visualized using UCSC Genome Browser to verify that DMRs were correctly annotated. BLASTx and the UCSC Genome Browser were used against the NCBI nonredundant database to assign genes to DMRs and to identify Gene Ontology (GO) terms. Genes with DMCs were input into g:Profiler for functional profiling [50].

We used ANCOVA (JMP 18.0.0) to determine if there was an association between either behavior and the mean percent methylation for five DMRs: *pkd2-l1*, *xiap*, *tbcb*, *ada2b*, and LOC103476631. These DMRs were chosen because of their significant differences in methylation between the two groups and because of the relevance of their biological function to the altered behaviors observed. Mean methylation percentage data were arcsine square-root transformed because they were percentages. Age at testing, testing date, and sequencing batch were included as covariates in the original model but were eliminated when p > 0.1. Sex and sibling group ID were included in the original model with the latter included as a random effect. Two- and three-way interactions were included in the original model and eliminated when p > 0.1. The assumptions of the test were met. The figure was plotted using a reduced major axis regression because it is formulated to handle errors in both the x and y variables.

## Data Availability

The raw data sequencing reads and metadata that support the findings (RNA-seq and RRBS reads) of this study are publicly available in the NCBI Sequence Read Archive under BioProject PRJNA1379867. Raw behavior data is provided in supplementary file S3.

## Ethics Statement

As outlined in our previous study using these fish (De Serrano et al, 2021), all procedures outlined in this study were approved by the Animal Care Committee at the University of Toronto (protocol numbers: 20008920, 20008921, 20009555, 20010160, 20010588, 20009045, 20010020, 20010527). Authorization to import and administer methylphenidate hydrochloride was obtained from the National Compliance and Exemption Division, Office of Controlled Substances, Health Canada (Authorization number: 26982.01.12). All experiments were performed in accordance with ARRIVE guidelines.

## Acknowledgements

We would like to acknowledge the National Council on Science and Engineering of Canada for support on this project (DG to IA, HR and NSERC CGSM to RJA). DJS is supported by the Ontario Genomics-CANSSI Postdoctoral Fellowship in Genome Data Science, the NSERC-PDF scholarship, and the McLaughlin Centre Scholars Grant.

## Supplemental Figure Captions

**Fig S1:**
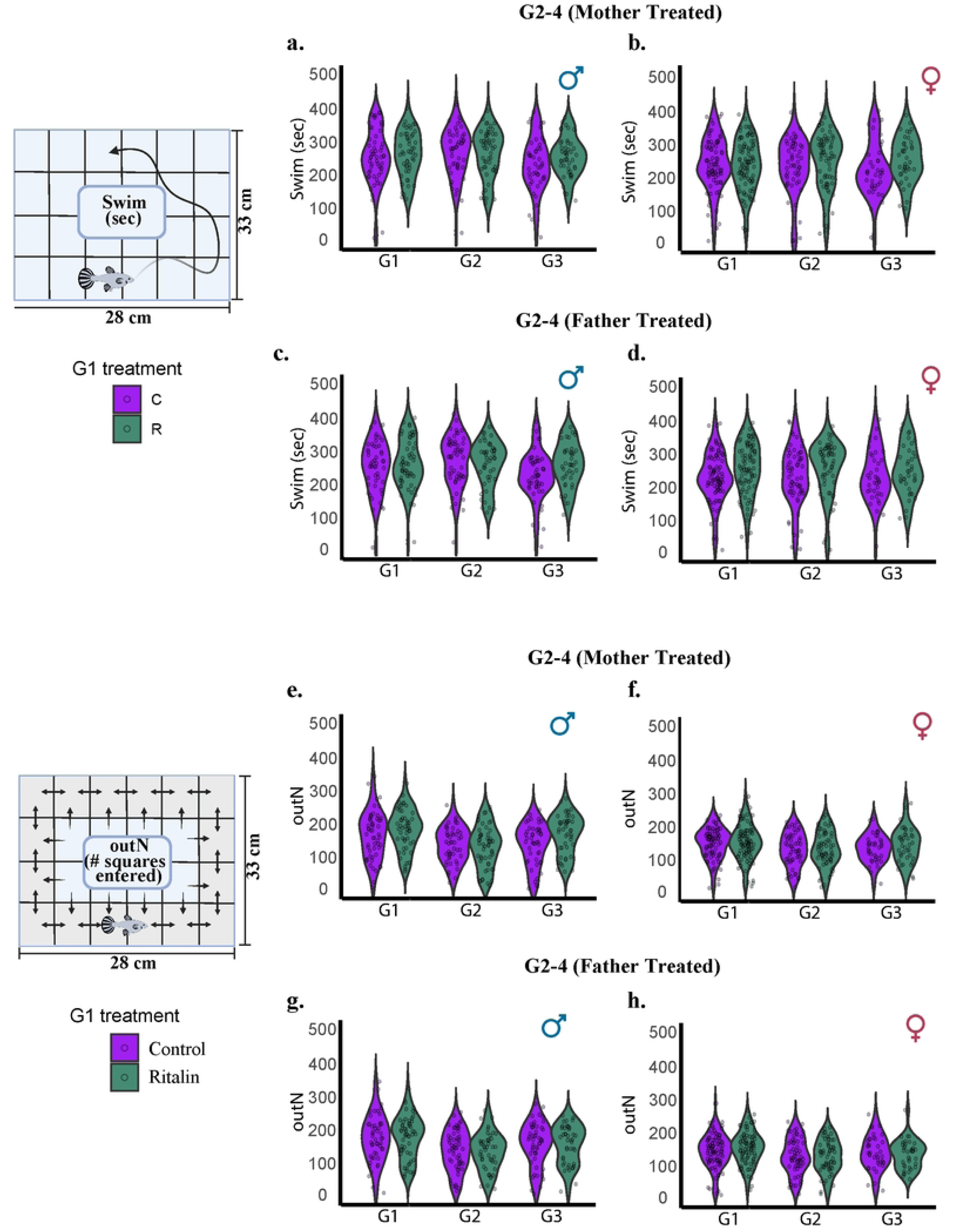
Effect of generation (G2-G4) on Swim and outer squares. **c-d.** There was a significant effect of generation on Swim for males and females with MPH treated fathers. There was a significant effect of generation on outer squares for males, but not females with MPH treated fathers. **c-d.** There was not a significant interaction between generation and G1 treatment for G2-G4 males.

**Fig S2:**
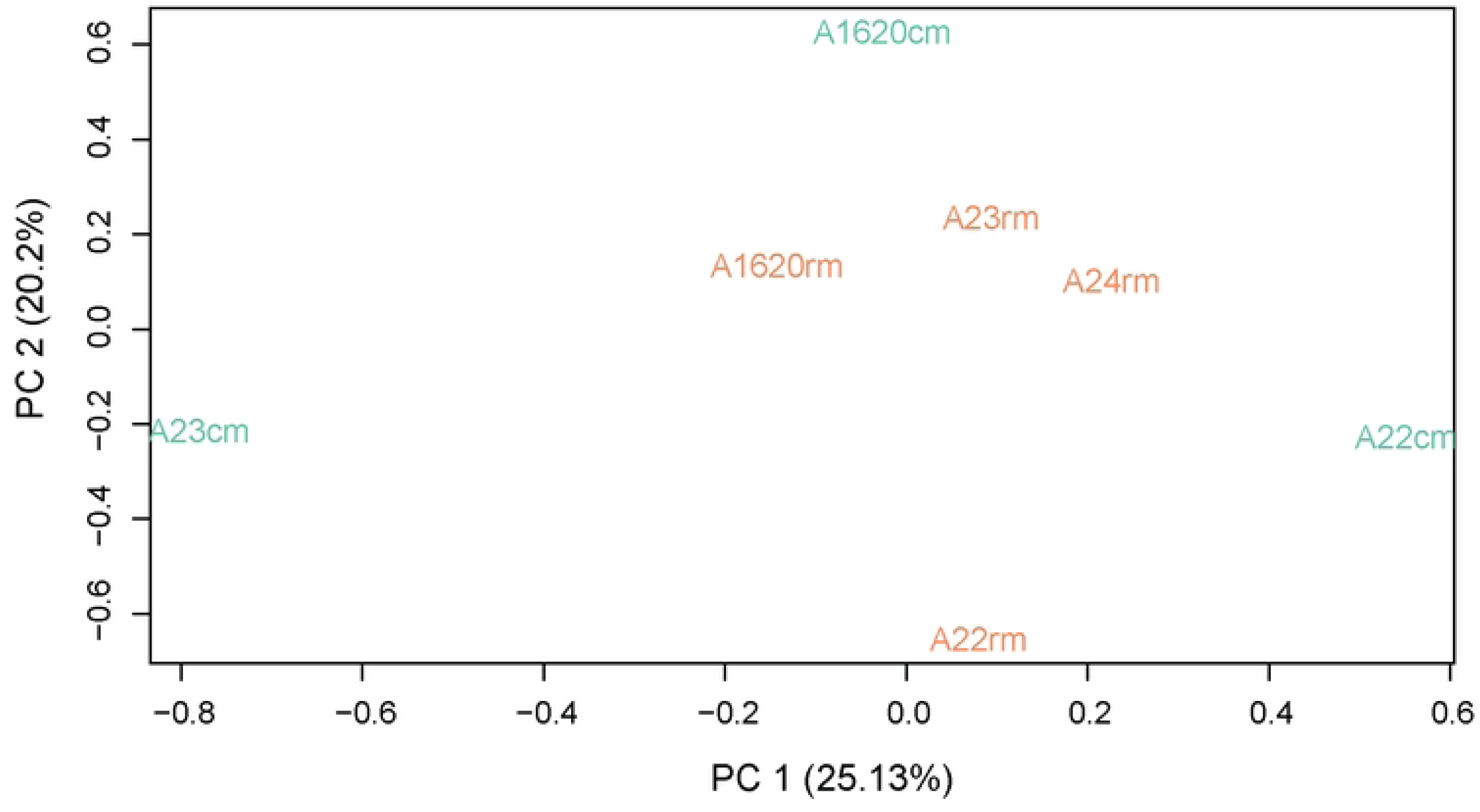
**a.** Principal component analysis of normalized RNA-Seq G_1_ samples. **b.** Principal component analysis of the genome-wide variation in methylation of brains of G4 fish.

**Fig S3:**
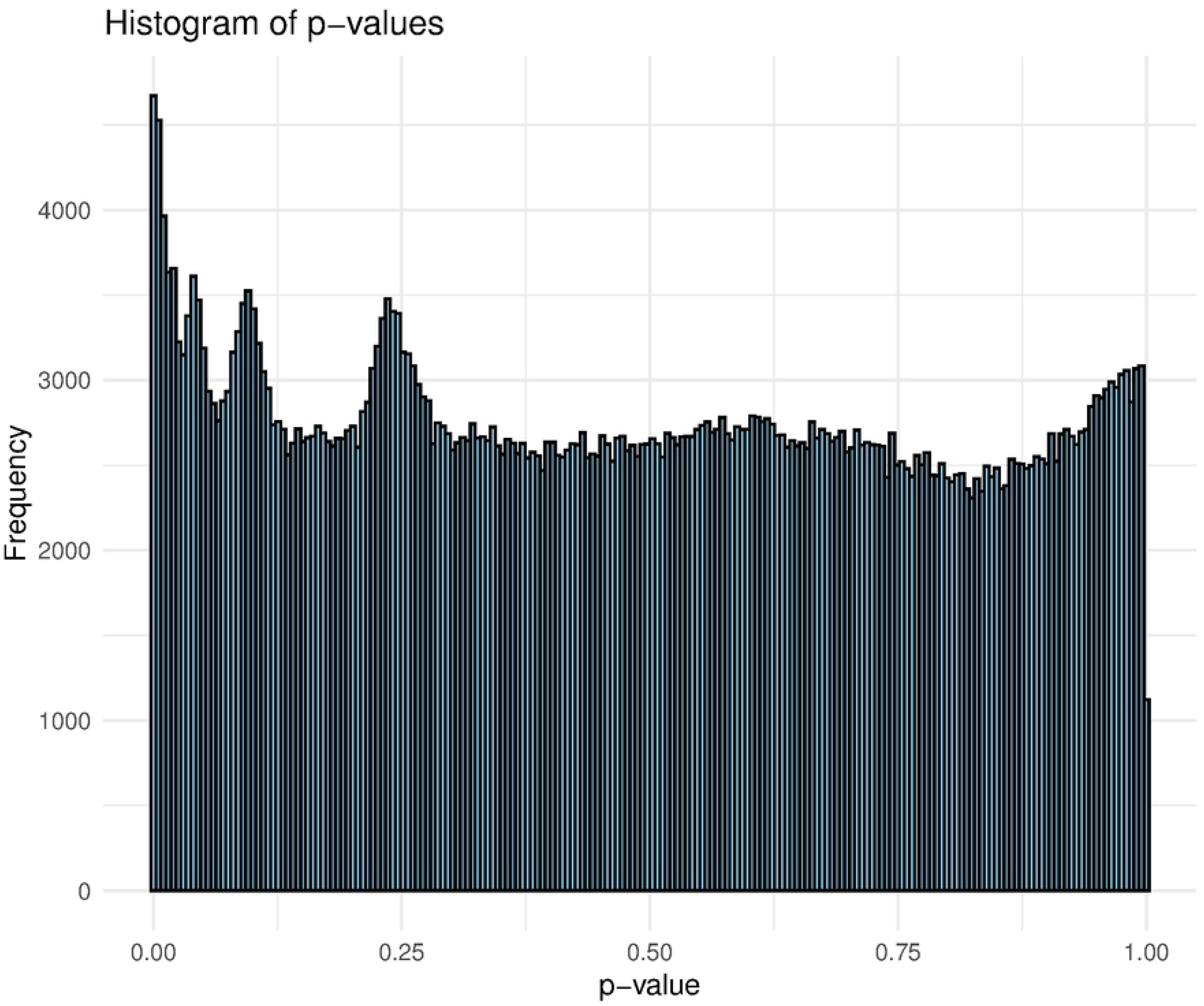
Histogram of p-value distribution for differentially methylated CPGs between G4 fish with MPH-treated ancestry and control.

## References

[1] Fitz-James MH, Cavalli G. Molecular mechanisms of transgenerational epigenetic inheritance. Nat Rev Genet. 2022 Jun;23(6):325–41.

[2] Nakao M. Epigenetics: interaction of DNA methylation and chromatin. Gene. 2001 Oct 31;278(1):25–31.

[3] Morgan HD, Sutherland HGE, Martin DIK, Whitelaw E. Epigenetic inheritance at the agouti locus in the mouse. Nat Genet. 1999 Nov;23(3):314–8.

[4] Weaver ICG, Cervoni N, Champagne FA, D’Alessio AC, Sharma S, Seckl JR, et al. Epigenetic programming by maternal behavior. Nat Neurosci. 2004 Aug;7(8):847–54.

[5] Keleher MR, Zaidi R, Hicks L, Shah S, Xing X, Li D, et al. A high-fat diet alters genome-wide DNA methylation and gene expression in SM/J mice. BMC Genomics. 2018 Dec 7;19(1):888.

[6] Lan X, Cretney EC, Kropp J, Khateeb K, Berg MA, Penagaricano F, et al. Maternal diet during pregnancy induces gene expression and DNA methylation changes in fetal tissues in sheep. Front Genet. 2013;4:49.

[7] Braz CU, Taylor T, Namous H, Rodriguez Lopez CM, Knight MI, Kelsey G, et al. Paternal diet induces transgenerational epigenetic inheritance of DNA methylation signatures and phenotypes in a sheep model. Proc Natl Acad Sci U S A. 2022;119(16):e2118096119.

[8] Dias BG, Ressler KJ. Parental olfactory experience influences behavior and neural structure in subsequent generations. Nat Neurosci. 2014;17(1):89–96.

[9] Anway MD, Cupp AS, Uzumcu M, Skinner MK. Epigenetic Transgenerational Actions of Endocrine Disruptors and Male Fertility. Science. 2005 Jun 3;308(5727):1466–9.

[10] Skinner MK, Ben Maamar M, Sadler-Riggleman I, Beck D, Nilsson E, McBirney M, et al. Alterations in sperm DNA methylation, non-coding RNA and histone retention associate with DDT-induced epigenetic transgenerational inheritance of disease. Epigenetics & Chromatin. 2018 Feb 27;11(1):8.

[11] Goldberg LR, Zeid D, Kutlu MG, Cole RD, Lallai V, Sebastian A, et al. Paternal nicotine enhances fear memory, reduces nicotine administration, and alters hippocampal genetic and neural function in offspring. Addiction Biology. 2021;26(1):e12859.

[12] Le Q, Yan B, Yu X, Li Y, Song H, Zhu H, et al. Drug-seeking motivation level in male rats determines offspring susceptibility or resistance to cocaine-seeking behaviour. Nat Commun. 2017 May 30;8(1):15527.

[13] Chan AYL, Ma TT, Lau WCY, Ip P, Coghill D, Gao L, et al. Attention-deficit/hyperactivity disorder medication consumption in 64 countries and regions from 2015 to 2019: a longitudinal study. EClinicalMedicine. 2023 Mar 20;58:101780.

[14] McCarthy S, Wilton L, Murray ML, Hodgkins P, Asherson P, Wong IC. Persistence of pharmacological treatment into adulthood, in UK primary care, for ADHD patients who started treatment in childhood or adolescence. BMC Psychiatry. 2012 Dec 5;12(1):219.

[15] Board AR, Guy G, Jones CM, Hoots B. Trends in stimulant dispensing by age, sex, state of residence, and prescriber specialty — United States, 2014–2019. Drug and Alcohol Dependence. 2020 Dec 1;217:108297.

[16] Quansah E, Ruiz-Rodado V, Grootveld M, Probert F, Zetterström TSC. 1H NMR-based metabolomics reveals neurochemical alterations in the brain of adolescent rats following acute methylphenidate administration. Neurochemistry International. 2017 Sep 1;108:109–20.

[17] Simchon-Tenenbaum Y, Weizman A, Rehavi M. Alterations in brain neurotrophic and glial factors following early age chronic methylphenidate and cocaine administration. Behavioural Brain Research. 2015 Apr 1;282:125–32.

[18] Bolaños CA, Barrot M, Berton O, Wallace-Black D, Nestler EJ. Methylphenidate treatment during pre- and periadolescence alters behavioral responses to emotional stimuli at adulthood. Biol Psychiatry. 2003;54(12):1317–1329.

[19] Montagnini BG, Mesquita SFP, Santos BD, Del Vecchio G, Perobelli JE, Kempinas WDG. Effects of repeated administration of methylphenidate on reproductive parameters of adult male rats. Physiol Behav. 2014;133:122–129.

[20] Csoka AB, Szyf M. Epigenetic side-effects of common pharmaceuticals: A potential new field in medicine and pharmacology. Medical Hypotheses. 2009 Nov 1;73(5):770–80.

[21] Hall ZJ, De Serrano AR, Rodd FH, Tropepe V. Casting a wider fish net on animal models in neuropsychiatric research. Progress in Neuro-Psychopharmacology and Biological Psychiatry. 2014 Dec 3;55:7–15.

[22] Hu J, Pérez-Jvostov F, Blondel L, Barrett RDH. Genome-wide DNA methylation signatures of infection status in Trinidadian guppies (Poecilia reticulata). Molecular Ecology. 2018;27(15):3087–102.

[23] Kelley JL, Tobler M, Beck D, Sadler-Riggleman I, Quackenbush CR, Arias Rodriguez L, et al. Epigenetic inheritance of DNA methylation changes in fish living in hydrogen sulfide–rich springs. Proceedings of the National Academy of Sciences. 2021 Jun 29;118(26):e2014929118.

[24] De Serrano AR, Hughes KA, Rodd FH. Paternal exposure to a common pharmaceutical (Ritalin) has transgenerational effects on the behaviour of Trinidadian guppies. Sci Rep. 2021 Feb 17;11(1):3985.

[25] Janowski BA, Huffman KE, Schwartz JC, Ram R, Hardy D, Shames DS, et al. Inhibiting gene expression at transcription start sites in chromosomal DNA with antigene RNAs. Nat Chem Biol. 2005 Sep;1(4):216–22.

[26] Yang Z, Xu F, Teschendorff AE, Zhao Y, Yao L, Li J, et al. Insights into the role of long non-coding RNAs in DNA methylation-mediated transcriptional regulation. Front Mol Biosci. 2022 Dec 2;9:1067406.

[27] Huang W, Li H, Yu Q, Xiao W, Wang DO. LncRNA-mediated DNA methylation: an emerging mechanism in cancer and beyond. J Exp Clin Cancer Res. 2022 Mar 15;41(1):100.

[28] Mohammad F, Pandey GK, Mondal T, Enroth S, Redrup L, Gyllensten U, et al. Long noncoding RNA-mediated maintenance of DNA methylation and transcriptional gene silencing. Development. 2012 Aug 1;139(15):2792–803.

[29] Zheng X, Li Z, Wang G, Wang H, Zhou Y, Zhao X, et al. Sperm epigenetic alterations contribute to inter- and transgenerational effects of paternal exposure to long-term psychological stress via evading offspring embryonic reprogramming. Cell Discov. 2021 Oct 27;7(1):1–22.

[30] Fouchécourt S, Picolo F, Elis S, Lécureuil C, Thélie A, Govoroun M, et al. An evolutionary approach to recover genes predominantly expressed in the testes of the zebrafish, chicken and mouse. BMC Evol Biol. 2019 Jul 3;19(1):137.

[31] Orts-Del’Immagine A, Seddik R, Tell F, Airault C, Er-Raoui G, Najimi M, et al. A single polycystic kidney disease 2-like 1 channel opening acts as a spike generator in cerebrospinal fluid-contacting neurons of adult mouse brainstem. Neuropharmacology. 2016 Feb 1;101:549–65.

[32] Djenoune L, Khabou H, Joubert F, Quan FB, Nunes Figueiredo S, Bodineau L, et al. Investigation of spinal cerebrospinal fluid-contacting neurons expressing PKD2L1: Evidence for a conserved system from fish to primates. Front Neuroanat. 2014;8:26.

[33] Delling M, DeCaen PG, Doerner JF, Febvay S, Clapham DE. Primary cilia are specialized calcium signalling organelles. Nature. 2013 Dec;504(7479):311–4.

[34] Böhm UL, Prendergast A, Djenoune L, Nunes Figueiredo S, Gomez J, Stokes C, et al. CSF-contacting neurons regulate locomotion by relaying mechanical stimuli to spinal circuits. Nat Commun. 2016 Mar 7;7(1):10866.

[35] Marie-Hardy L, Slimani L, Messa G, El Bourakkadi Z, Prigent A, Sayetta C, et al. Loss of CSF-contacting neuron sensory function is associated with a hyper-kyphosis of the spine reminiscent of Scheuermann’s disease. Sci Rep. 2023 Apr 4;13(1):5529.

[36] Rosemberg DB, Rico EP, Senger MR, Dias RD, Bogo MR, Bonan CD, Souza DO. Kinetic characterization of adenosine deaminase activity in zebrafish (Danio rerio) brain. Comp Biochem Physiol B Biochem Mol Biol. 2008 Sep;151(1):96–101.

[37] Dale N. Delayed production of adenosine underlies temporal modulation of swimming in frog embryo. J Physiol. 1998 Aug 15;511 (Pt 1)(Pt 1):265–72.

[38] Chen H, Xing G, Xu W, Chen Y, Xia L, Huang H, et al. The adenosine A2A receptor in human sperm: its role in sperm motility and association with in vitro fertilization outcomes. Front Endocrinol (Lausanne). 2024 May 30;15:1410370.

[39] Boehmler W, Petko J, Woll M, Frey C, Thisse B, Thisse C, et al. Identification of zebrafish A2 adenosine receptors and expression in developing embryos. Gene Expr Patterns. 2009 Mar;9(3):144–51.

[40] Dharmadhikari AV, Kang SH, Szafranski P, Person RE, Sampath S, Prakash SK, et al. Small rare recurrent deletions and reciprocal duplications in 2q21.1, including brain-specific ARHGEF4 and GPR148. Hum Mol Genet. 2012 Aug 1;21(15):3345–55.

[41] Querne L, Fall S, Le Moing AG, Bourel-Ponchel E, Delignières A, Simonnot A, et al. Effects of Methylphenidate on Default-Mode Network/Task-Positive Network Synchronization in Children With ADHD. J Atten Disord. 2017 Dec;21(14):1208–1220.

[42] Chan JYC, Dennis TA, MacLeod MA. The over-prescription of Ritalin for suspected cases of ADHD [Internet]. Ottawa (ON): University of Ottawa; 2012 Jan [cited 2024 May 31]. Available from: http://hdl.handle.net/10393/34386

[43] Wickham H. Getting started with ggplot2. In: Wickham H, editor. ggplot2: Elegant graphics for data analysis. 2nd ed. Cham: Springer International Publishing; 2016. p. 11–31.

[44] Andrews S. 2010. FastQC: a quality control tool for high throughput sequence data [Internet]. Cambridge (UK): Babraham Bioinformatics, Babraham Institute. Available from: https://www.bioinformatics.babraham.ac.uk/projects/fastqc

[45] Bolger AM, Lohse M, Usadel B. Trimmomatic: a flexible trimmer for Illumina sequence data. Bioinformatics. 2014 Aug 1;30(15):2114–20.

[46] Kim D, Paggi JM, Park C, Bennett C, Salzberg SL. Graph-based genome alignment and genotyping with HISAT2 and HISAT-genotype. Nat Biotechnol. 2019 Aug;37(8):907–15.

[47] Pertea M, Pertea GM, Antonescu CM, Chang TC, Mendell JT, Salzberg SL. StringTie enables improved reconstruction of a transcriptome from RNA-seq reads. Nat Biotechnol. 2015 Mar;33(3):290–5.

[48] Robinson MD, McCarthy DJ, Smyth GK. edgeR: a Bioconductor package for differential expression analysis of digital gene expression data. Bioinformatics. 2010 Jan 1;26(1):139–40.

[49] Risso D, Ngai J, Speed TP, Dudoit S. Normalization of RNA-seq data using factor analysis of control genes or samples. Nat Biotechnol. 2014 Sep;32(9):896–902.

[50] Kolberg L, Raudvere U, Kuzmin I, Adler P, Vilo J, Peterson H. g:Profiler—interoperable web service for functional enrichment analysis and gene identifier mapping (2023 update). Nucleic Acids Research. 2023 Jul 5;51(W1):W207–12.

[51] Krueger F. Trim Galore: A wrapper around Cutadapt and FastQC to consistently apply adapter and quality trimming to FastQ files. 2012

[52] Krueger F, Andrews SR. Bismark: a flexible aligner and methylation caller for Bisulfite-Seq applications. Bioinformatics. 2011 Jun 1;27(11):1571–2.

[53] Breheny P, Stromberg A, Lambert J. p-Value Histograms: Inference and Diagnostics. High Throughput. 2018 Aug 31;7(3):23.

[54] Akalin A, Kormaksson M, Li S, Garrett-Bakelman FE, Figueroa ME, Melnick A, et al. methylKit: a comprehensive R package for the analysis of genome-wide DNA methylation profiles. Genome Biology. 2012 Oct 3;13(10):R87.

[55] Li S, Garrett-Bakelman FE, Akalin A, Zumbo P, Levine R, To BL, et al. An optimized algorithm for detecting and annotating regional differential methylation. BMC Bioinformatics. 2013 Apr 10;14(5):S10.

[56] Akalin A, Franke V, Vlahoviček K, Mason CE, Schübeler D. genomation: a toolkit to summarize, annotate and visualize genomic intervals. Bioinformatics. 2015 Apr 1;31(7):1127–9.

